# Co-occurring eQTLs and mQTLs: detecting shared causal variants and shared biological mechanisms

**DOI:** 10.1101/094656

**Authors:** Brandon L. Pierce, Lin Tong, Maria Argos, Farzana Jasmine, Muhammad Rakibuz-Zaman, Golam Sarwar, Md. Tariqul Islam, Hasan Shahriar, Tariqul Islam, Mahfuzar Rahman, Md. Yunus, Muhammad G. Kibriya, Lin S. Chen, Habibul Ahsan

**Affiliations:** Department of Public Health Sciences, The University of Chicago, Chicago, IL 60637; Department of Human Genetics, The University of Chicago, Chicago, IL 60637; Comprehensive Cancer Center, The University of Chicago, Chicago, IL 60637; UChicago Research Bangladesh, Mohakhali, Dhaka, Bangladesh; Research and Evaluation Division, BRAC, Dhaka, Bangladesh; International Centre for Diarrhoeal Disease Research, Bangladesh, Dhaka, Bangladesh; Department of Medicine, The University of Chicago, Chicago, IL 60637

**Keywords:** expression quantitative trait loci (eQTL), methylation quantitative trait loci (mQTL), DNA methylation, co-localization, mediation, partial correlation

## Abstract

Inherited genetic variation impacts local gene expression and DNA methylation in humans. Expression and methylation quantitative trait loci (cis-eQTLs and cis-mQTLs) often occur at the same genomic location, suggesting a common causal variant and shared mechanism. Using DNA and RNA from peripheral blood of Bangladeshi individuals, we use “co-localization” methods to identify 3,695 eQTL-mQTL pairs that are likely to share a causal variant. Using partial correlation analysis and mediation analysis, we identify >500 pairs with evidence of a causal relationships between expression and methylation (i.e., shared mechanism) with many additional pairs that we are underpowered to detect. These co-localized pairs are enriched for SNPs showing opposite effects on expression and methylation, although a many affect multiple CpGs in opposite directions. Evidence of shared SNP-age interaction also supports shared mechanisms for two eQTL-mQTL pairs. This work demonstrates the pervasiveness of co-regulated expression and methylation traits in the human genome. This approach can be applied to other types of molecular QTLs to enhance our understanding of regulatory mechanisms.

## Introduction

Genetic variation has a substantial impact on mRNA abundance in humans ^1^. Genome-wide scans to identify regions that harbor such variants, regions known as eQTLs (expression quantitative trait loci), have been conducted using RNA from a wide-array of human tissue types and cell types, and eQTLs have been identified for the vast majority of human genes.

In addition to studies of transcript abundance, recent work has described the effects of genetic variation on other genomic and cellular phenotypes, such as DNA methylation ^2–5^, DNase hypersensitivity ^6^, histone modifications and nucleosome positioning^7^, RNA splicing^8,9^, translational efficiency/ribosome occupancy^10,11^, and protein abundance ^12,13^. Because many QTLs appear to influence multiple local molecular phenotypes, there is great interest in identifying variants that have coordinated effects on multiple phenotypes and understanding the mechanisms by which such variants act.

Recently, several groups have identified SNPs associated with both expression of nearby genes and methylation of nearby CpG sites ^14–17^. For these cis-eQTLs that also appear to be local methylation-QTL (cis-mQTLs), it is possible that co-occurring eQTLs and mQTLs share a common causal variant, suggesting a shared biological mechanism by which the causal variant influences both expression and methylation. methylation could be reactive to expression (i.e., methylation responds to genetically-determined variation in gene expression, perhaps due to a SNP’s effect on transcription factor binding), or methylation could mediate the effect of the SNP on expression (i.e., increased promoter methylation suppresses transcription factor binding). For such co-occurring eQTLs and mQTLs, several groups have developed and applied approaches intended to determine if a causal relationship exists between the local DNA methylation and expression, including likelihood-based approaches^17^, Bayesian network approaches ^16^, and partial correlation approaches ^18^.

One limitation of the prior work on this topic is a lack of rigorous assessment of the hypothesis that co-occurring eQTLs and mQTLs share a common causal variant. Recently developed tests for “colocalization” allow one to assess whether two association signals are consistent with a shared causal variant^19^. Using summary statistics for an eQTL and an mQTL, one can estimate the probability that the eQTL and mQTL share a common causal variant. This information can be used to guide subsequent studies of co-occurring eQTL/mQTL pairs.

In this work, we use genome-wide data on SNPs and array-based expression and DNA methylation from South Asian individuals to identify cis-eQTLs and cis-mQTLs. We describe the extent to which the observed cis-eQTLs and cis-mQTLs share common causal variants using co-localization methods ^19^. Using eQTL/mQTL pairs with a high probability of sharing a causal variant, we then assess the evidence that expression and methylation are causally related to one another using partial correlation analysis and mediation analysis. In addition, we characterize these co-localized QTLs and search for SNP-age and SNP-sex interactions that are shared across co-localized QTLs.

## Results

A simple overview of our workflow for identifying eQTLs and mQTLs that are likely to share a common causal variant (CCV) is shown in **Figure 1**. A more detailed workflow is provided in **Supplementary Figure 1**.

**Figure 1.**
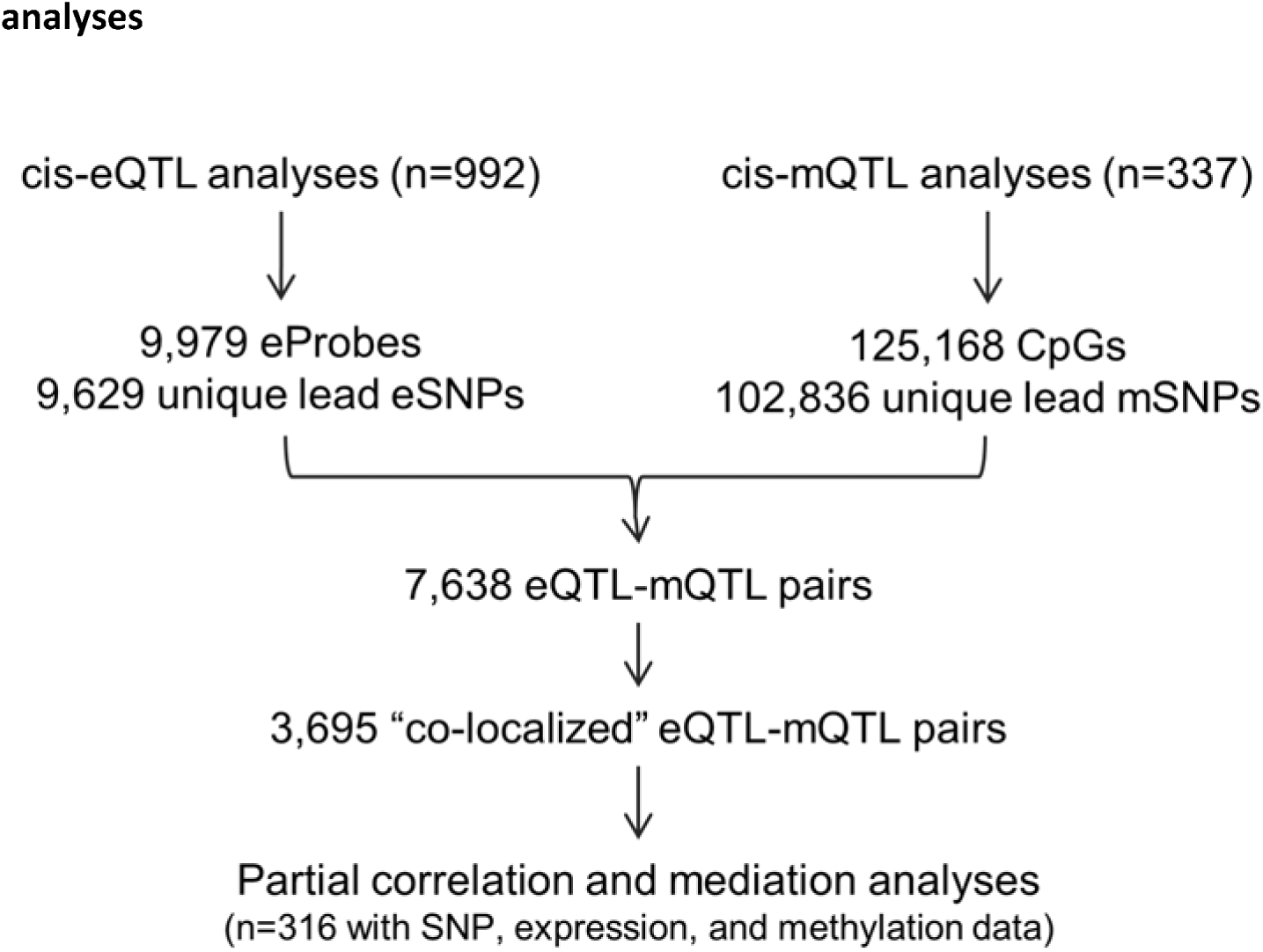
Summary of the workflow for cis-QTL, co-localiation, partial correlation, and mediation analyses.

### Observed cis-eQTLS and cis-mQTLs

We conducted genome-wide eQTL and mQTL analyses using data on 992 and 337 non-overlapping study participants, respectively. Patterns of peripheral blood cis-eQTLs and cis-mQTLs are reported in **Table 1**. At an FDR of 0.01, we detected a *cis*-eQTL for 9,979 expression probes, corresponding to 8,115 genes (i.e.cis-eGenes), and 9,629 unique lead eSNPs. At an FDR of 0.01, we detected evidence of an mQTL for 125,162 CpG sites, corresponding to 102,836 unique lead mSNPs. 90,709 of these CpG sites were assigned to a total of 16,227 genes (based on llluminás annotation). On average, lead mSNPs were 10 kb closer to their target CpGs than lead eSNPs were to their target transcription start site (p=10^−19^, controlling for QTL P-value).

**Table 1.**
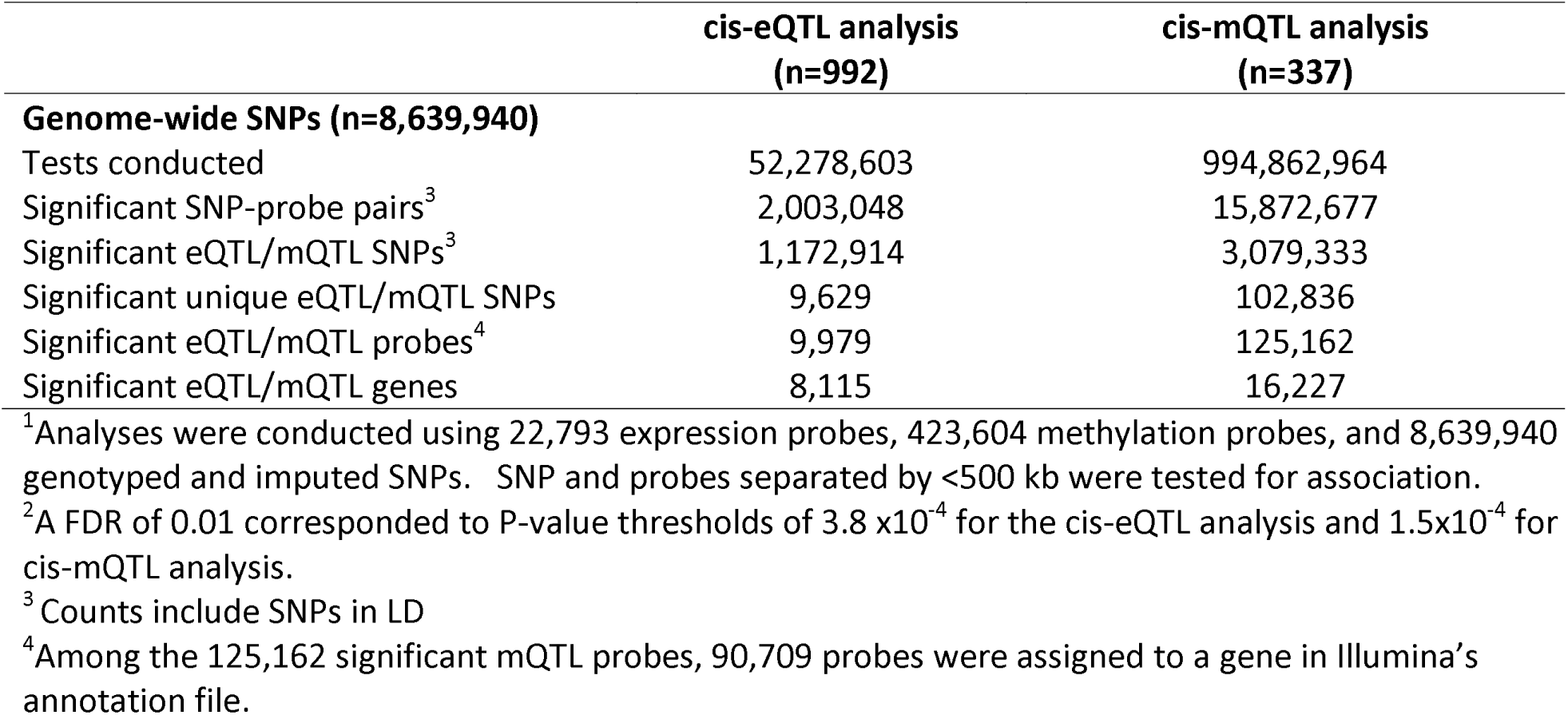
Summary of cis-eQTL and cis-mQTL signals identified in genome-wide^1^ analyses using a false discovery rate (FDR^2^) of 0.01

### Co-localization of cis-eQTLs and cis-mQTLs

7,454 of our 9,629 unique eSNPs were among the mSNPs identified at an FDR of 0.01, corresponding to 7,638 unique SNP-eProbe-CpG combinations potentially representing a CCV. Using these pairs of eProbes and CpGs associated with a common SNP, we conducted a Bayesian test of co-localization ^19^ for each of the 7,638 pairs (see methods) (**Supplementary Table 1**). This method assesses whether two association s signals are consistent with a CCV. A substantial number of these test (~20%) produced probabilities of CCV that were very close to zero, but probability of CCV was strongly related to the linkage disequilibrium (LD) between the lead eSNP and the lead mSNP, with low LD corresponding to low probability of CCV (**Figure 2A**). Removing eProbe-CpG pairs whose lead eSNPs and mSNPs were in weak LD (r^2^<0.5) eliminated the majority of the eProbe-CpG pairs showing very weak evidence of co-localization (**Figure 2B**). These results reflect the existence to two major types of pairs: those very likely to share a CCV and those that do not share a CCV.

**Figure 2.**
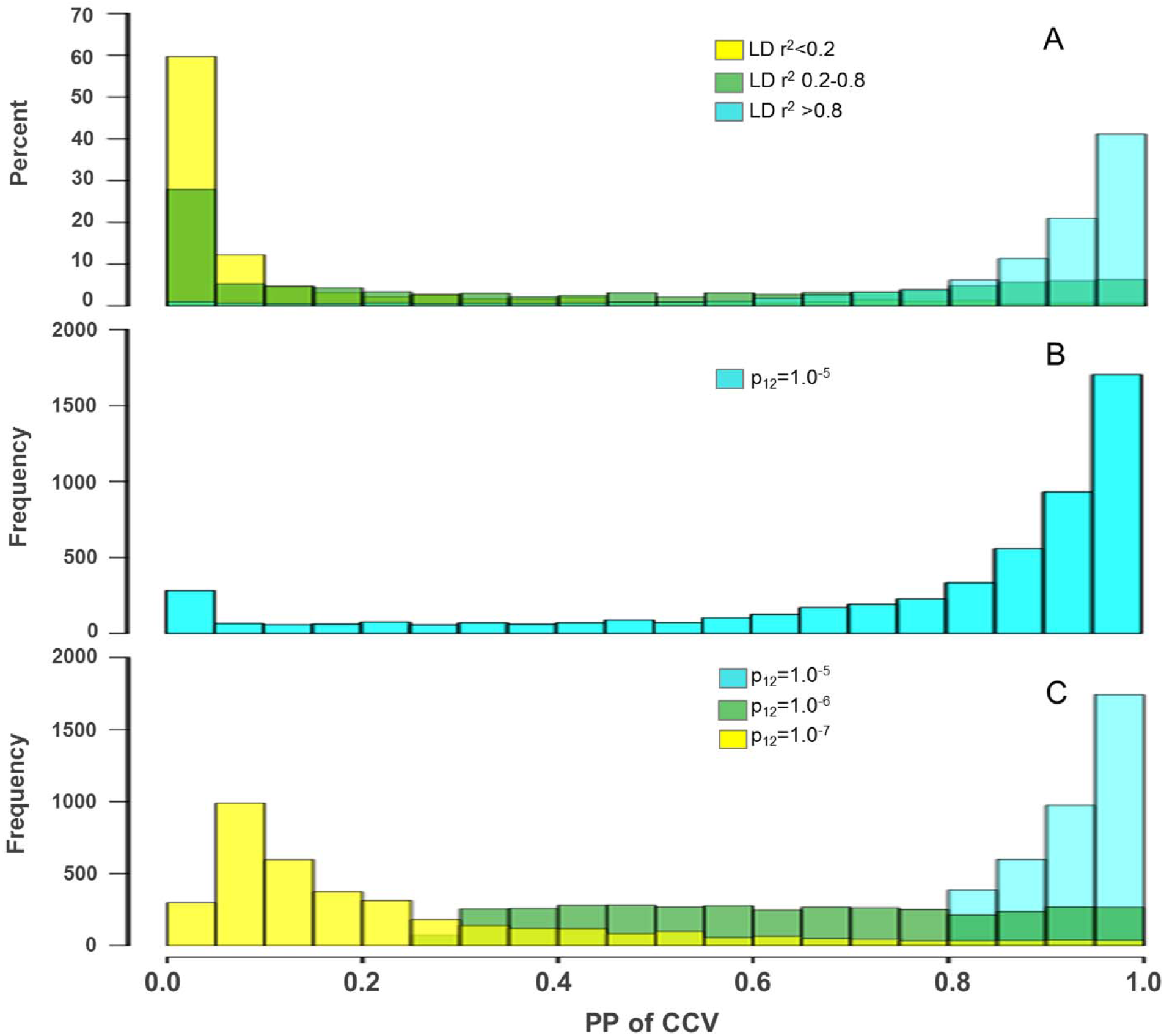
The distribution of the posterior probability (PP) of a common causal variant (CCV) for cooccurring eQTL-mQTL pairs. A: PP of a CCV at different values of LD (r^2^) between the lead eSNP and lead mSNP for n=7,638 eQTL-mQTL pairs. B: 5,286 eQTL-mQTL pairs with r^2^ >0.5 between lead eSNP and lead mSNP. C: 3,695 eQTL-mQTL pairs with a PP of a CCV >80% for p_12_=10^−5^. The distribution of PP of CCV for these pairs is also shown at p_12_ values of 10^−6^ and 10^−7^.

### Sensitivity of the probability of CCV to the choice of prior

The Bayesian co-localization ^19^ requires specifying a prior probability for a SNP being associated with expression (p_1_), methylation (p_2_), and both traits (p_12_). Following the approach used by the method developers ^19^, we used a prior of 10^−4^ for p_1_ and p_2_, and we varied the value of p_12_ (10^−5^, 10^−6^, and 10^−7^). Among all 7,638 pairs, we observed 3,695 pairs with a probability of CCV >80%, and we designated these as potentially “co-localized” eQTL-mQTL pairs (based on p_12_ = 10^−5^). We selected these for further analysis (**Supplementary Table 1**). The majority of co-localization tests were highly sensitive to p_12_ (**Figure 2C**), and these priors are interpreted respectively as 1 in 10, 1 in 100, and 1 in 1000 probability that a SNP associated with expression is also associated with methylation (or vice versa) given value of 10^−4^ for both pi and p_2_ (see Methods). Because decreasing p_12_ to 10^−7^ eliminates the vast majority of evidence for co-localization, we used the threshold of probability of a CCV >0.80 at p_12_ = 10^−5^ to define potentially co-localized pairs. Six of our strongest co-localized signals (based on probability of a CCV) are shown in **Figure 3**.

**Figure 3.**
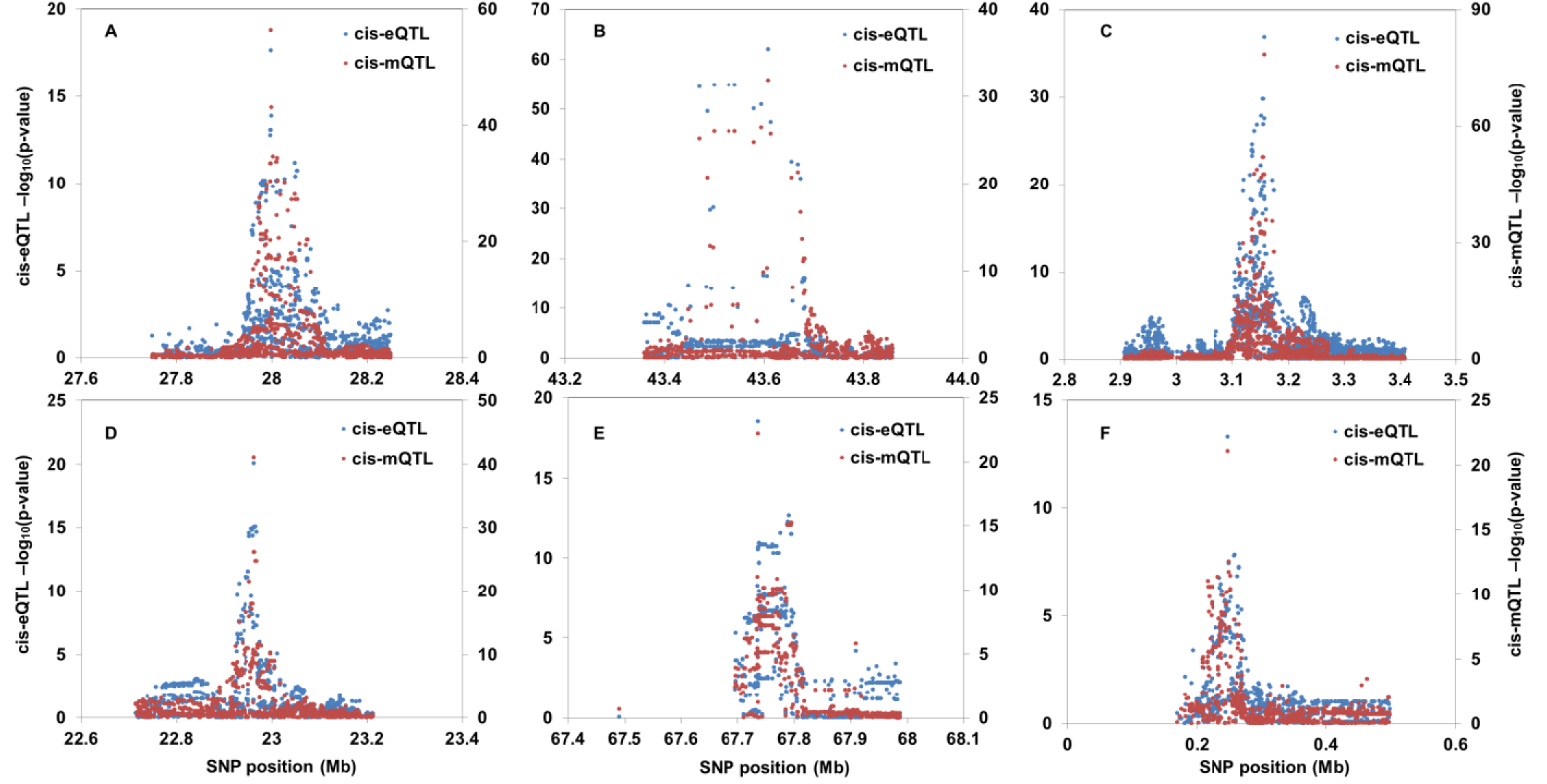
Examples of 6 co-localized eQTL-mQTL pairs. A: ILMN_1658464 (GTF3A) and cg22138327. B: ILMN_1694711 (MAD2L1BP) and cg14302083. C: ILMN_1721978 (CARD11) and cg19214707. D: ILMN_1737918 (C1QA) and cg10916651. E: ILMN_2193591 (UNC93B1) and cg20272935. F: ILMN_2282366 (IQSEC3)and cg25396728.

### Co-localization and LD

The probability of a CCV showed a strong inverse association with the “LD score” of the lead SNP(**Figure 4A and Supplementary Table 2**) with LD score defined as the sum of the pairwise r^2^ values between the LD SNP and all SNPs within 500 kb^20^. This result demonstrates the increased uncertainty regarding sharing of a causal variant in regions containing many highly correlated variants. Co-localized eQTL-mQTL pairs with high LD scores for the lead SNPs also appear to have probabilities of CCV values that were more sensitive to the prior than pairs with low LD scores (**Figure 4B** **and** **4C**), indicating that the test for co-localization is better able to detect evidence of a shared causal in regions of low LD. The probability of CCV was also strongly associated with the strength of the eQTL and mQTL association signals (in terms of P-value) (**Supplementary Table 2**).

**Figure 4.**
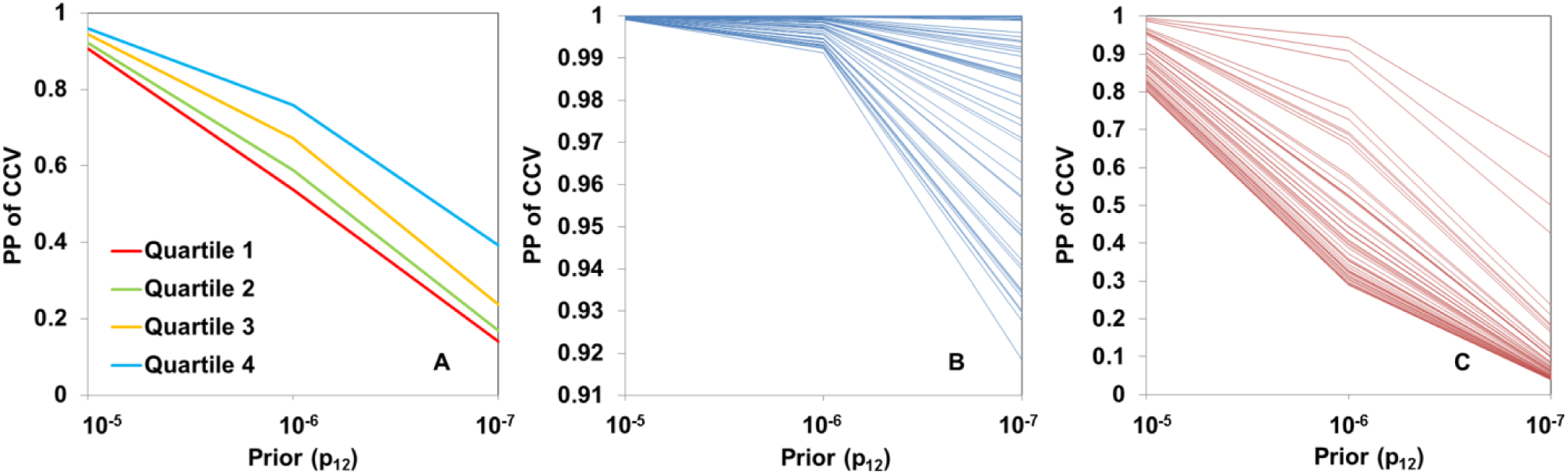
The posterior probability (PP) of sharing a common causal variant (CCV) depends on the extent of LD in the region. A: Average PPs of CCV at different prior (p_12_) by lead SNP stratified by LD score for 3,695 SNPs. B: PP of CCV at different prior (p_12_) for the lead SNPs with the lowest LD score (50 SNPs). C: PP of CCV at different prior (p_12_) for the 50 lead SNPs with the highest LD score (50 SNPs)

### Partial Correlation Analysis

For the 3,695 potentially co-localized pairs, we restricted analyses to 316 genotyped individuals with both expression and methylation data and conducted partial mediation analysis to determine if there was residual correlation between the expression probe and the CpG probe after removing the effects of the lead eQTL SNP (i.e., regressing the probe on the SNP and taking the residual). When a SNP has independent effects on expression and methylation, i.e., no causal correlation between expression and methylation, the residuals after removing lead SNP effects would be uncorrelated. As such, observing correlations in residuals provides support for a causal relationship between expression and methylation ^18^. We observed 507 eProbe-CpG pairs for which the correlation between the two remained significant (P<0.05) after adjustment for the lead SNP. Correlations tended to be weaker in magnitude after SNP-adjustment, and significant correlations were more likely to be negative than positive (**Figure 5A**).

**Figure 5.**
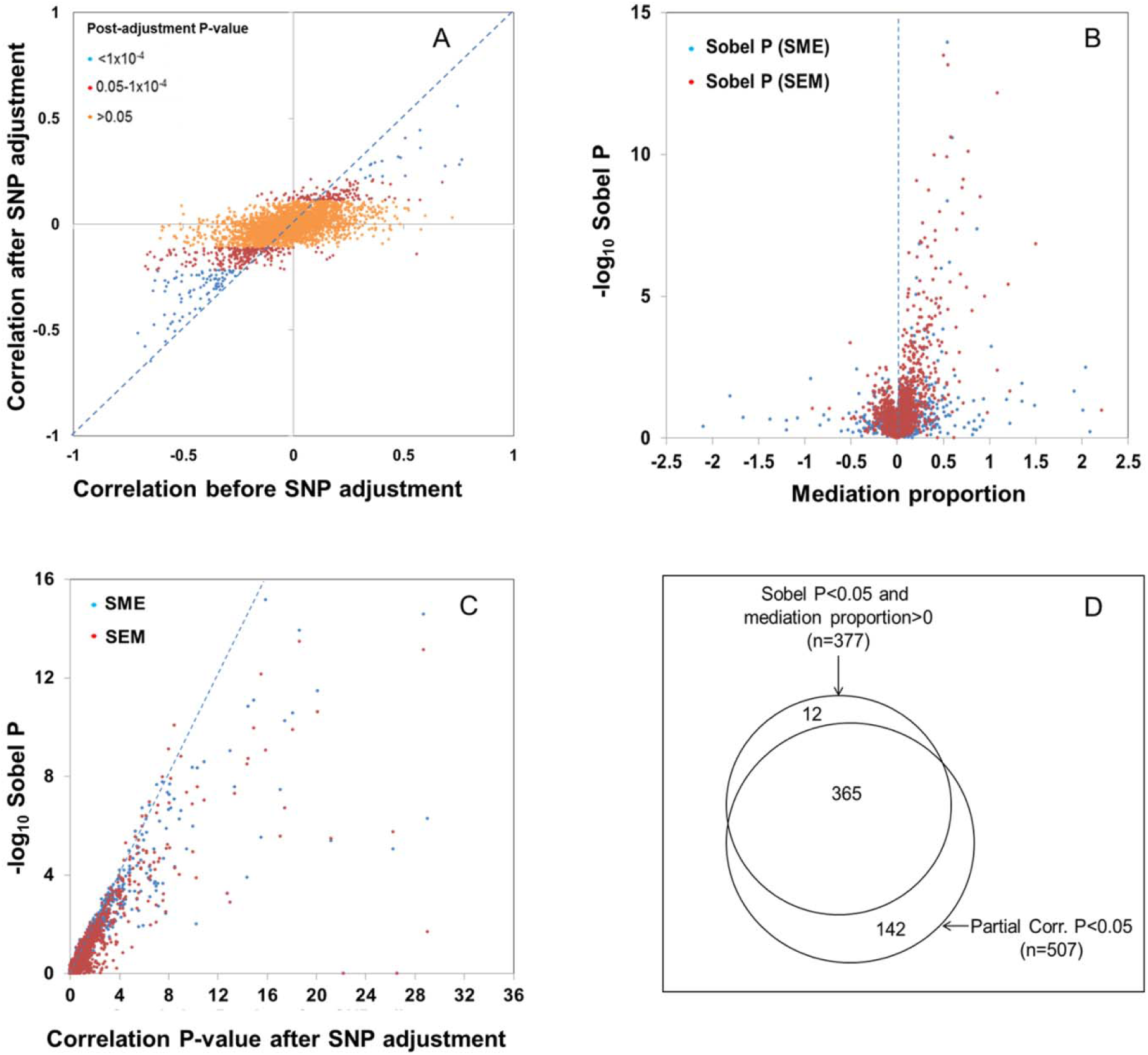
Partial correlation analysis and mediation analysis for 3,695 eQTL-mQTL pairs provides evidence for shared regulatory mechanisms. A: Partial correlation analysis results. B: Mediation analysis results for the SME and the SEM model. C: relationship P-value for mediation (Sobel P) and the post-adjustment correlation P-value from partial correlation analysis. D: Venn diagram showing the concordance between mediation and partial correlation analysis. All models includes adjustments for age, sex, and PCs from both the expression and methylation data (n=316).

### Mediation analysis

We applied mediation analysis to our 3,695 co-localized pairs as previously described ^21^ in order to assess evidence that 1) local DNA methylation mediates the effect of the SNP on local gene expression (SNP -> methylation -> Expression or “SME”) and 2) gene expression mediates the effect of the SNP on local DNA methylation (SNP -> Expression -> methylation or “EM”), a scenario in which DNA methylation is reactive to variation in gene expression activity. We observe significant evidence of mediation (Sobel P<0.05 *and %* mediation >0 for SEM *or* SME) for 377 pairs (**Figure 5B**). Evidence for mediation was often detected for specific gene-CpG pairs regardless of which mediation model was tested (SEM and SME, respectively) (**Supplementary Figure 2A**) demonstrating that mediation analysis is essentially a test for shared variance among the SNP, CpG, and expression trait and can be inadequate by itself for determining the direction of causality between two variables. However, we demonstrate using simulated data that evidence for mediation should be stronger when the causal model is correctly specified (**Supplementary Figure 3**).

### Comparison of Partial Correlation and Mediation Results

The mediation analysis results were highly consistent with the partial correlation analysis results, with nearly all of the 365 “mediated” pairs being among the 507 pairs with a significant partial correlation after SNP adjustment (**Figure 5D**). The distribution of the Sobel P-value and the post-adjustment P-value were similar, in that pairs with small Sobel P values tended to have small P-value for correlation after SNP adjustment (**Figures 5C**). Our ability to detect significant evidence of mediation and partial correlation was strongly related to the strength of the eQTL and mQTL being tested (**Supplementary Table 3**), indicating that we are likely underpowered to detect mediation and/or partial correlation for a substantial number to eQTL-mQTL pairs. In addition to partial correlation and mediation analyses, we also conducted Bayesian network analysis (BNA) as previously described ^16^ to determine if either the SME or SEM models were more strongly supported by the data than a model based on independent SNP effects on expression and methylation (see methods). BNA identified SEM or SME to be the most likely model for 572 pairs with >200 of the pairs also identified by either partial correlation and/or mediation analysis (**Supplementary Figure 2B and 2C**). For, the vast majority of these pairs (>500), SME was the most likely model, which was largely consistent with results from mediation analysis (**Supplementary Figure 2B**). Some consistency is expected as both methods rely on conditional dependence.

### Co-localized eQTL/mQTL pairs tend to of have opposite effects

Among our 3,695 co-localized eProbe-CpG pairs, the direction of the effect of the SNP on expression and methylation was more often in opposite directions (n=2,138; 57.8%) than in the same direction (n=1,557; 42.2%) (**Figure 6A**), consistent with the hypothesis that reduced promoter methylation is indicative of a more open chromatin state and increased transcriptional activity. When restricting to pairs that show evidence of a shared mechanism, according to either partial correlation or mediation (at either P<0.05 or P<0.001), a more striking difference is observed, with 70–80% of co-localized eQTL-mQTL pairs showing opposite directions of association, depending on the P-value threshold used (**Figure 6A**). Similarly, the expression and methylation traits for co-localized pairs were more often negatively correlated than positively correlated (57% negative). This imbalance was much stronger after restricting to pairs showing evidence of a shared causal mechanism, according to either partial correlation or mediation analysis, with 70–80% of co-localized eQTL-mQTL pairs showing an inverse correlation (based on P<0.05 and P<0.0001) (**Figure 6B**).

**Figure 6.**
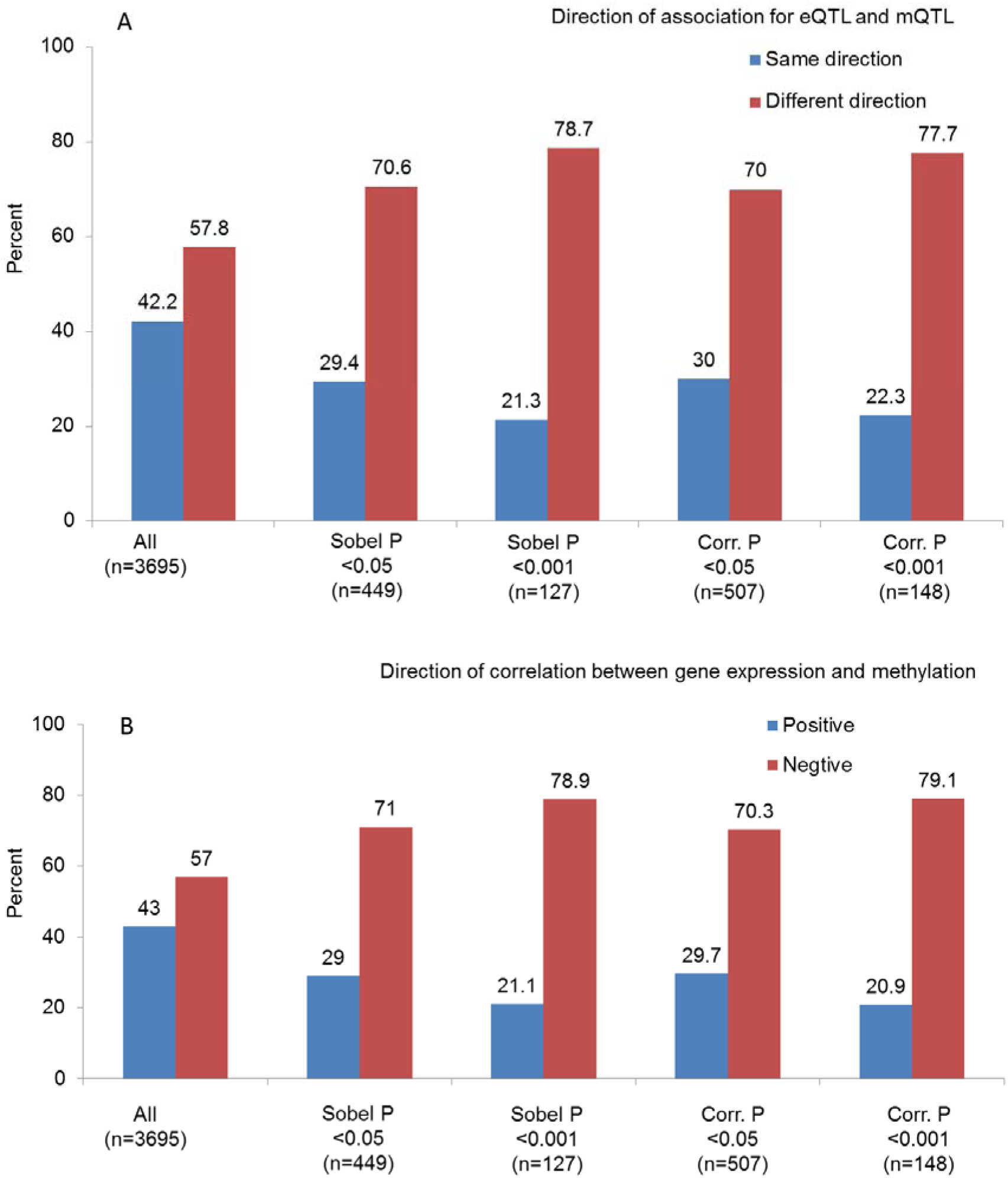
Proportion of potentially co-localized eQTL-mQTL pairs for which the SNP's association with expression and methylation is in the same or different directions. Overall results are shown, as well as those stratified by P-values from mediation analysis (Sobel P-value) and partial correlation analysis (post-adjustment P-value).

### eSNPs often co-localize with mSNPs associated with both increased and decreased methylation

There were 1,557 co-localized eQTL-mQTLs pairs showing associations with expression and methylation in the same direction (i.e., an allele is associated with an increase in both expression and methylation). Among these mQTLs, we searched for additional nearby CpG sites that showed an association with the SNP that was in the opposite direction of the eQTL. In 1,219 out of 1,557 cases, we identified at least one such a secondary CpG, and these CpGs were consistently inversely associated with the CpG originally selected. In other words, many of our eSNPs/mSNPs were associated with methylation at multiple nearby CpGs, with the minor allele increasing methylation at one CpG while decreasing methylation at another. The three examples with the strongest association between SNP and secondary CpG are shown in **Figure 7**, and all of these secondary mQTL signals also co-localize with the primary eQTL with probabilities of CCV > 90%.

**Figure 7.**
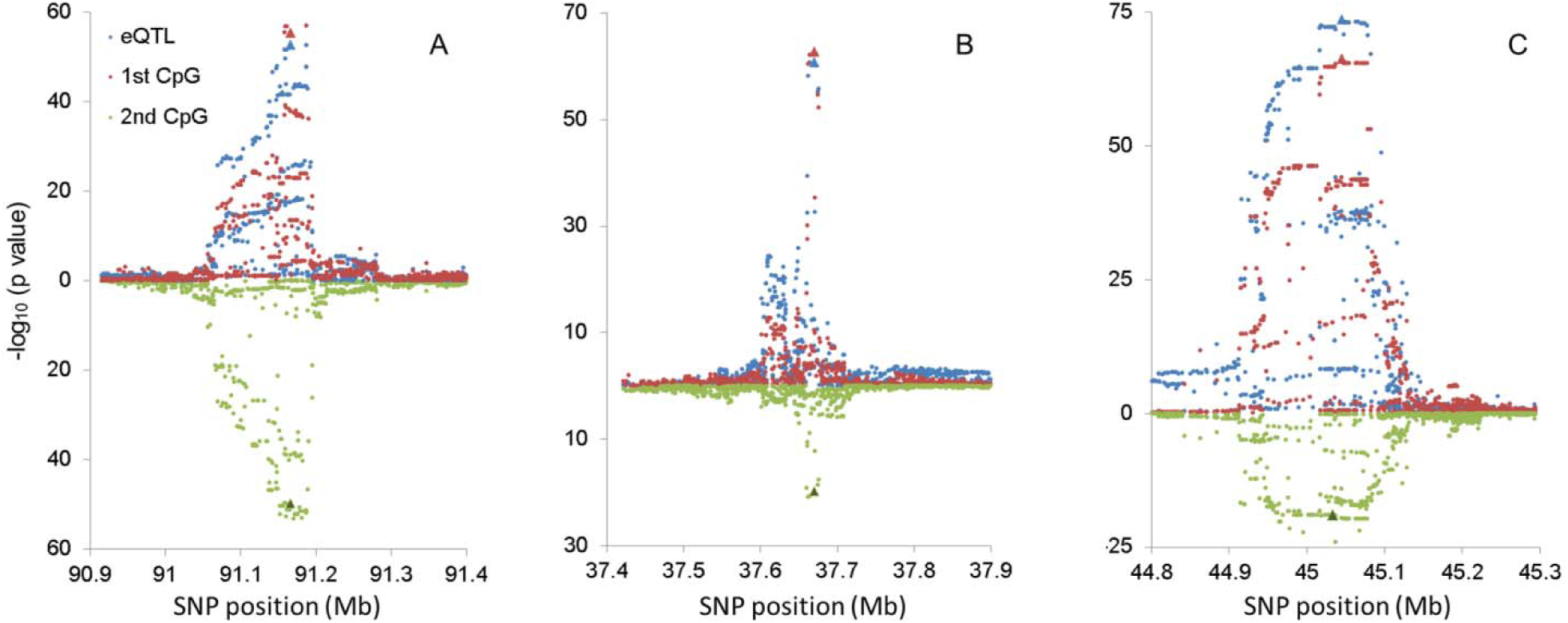
Examples of eQTLs which co-localize with an mQTL that has opposite effects on two nearby CpG sites. Results for the mQTL (red) and eQTL (blue) selected for co-localization analysis are shown above (ascending). For these scenarios, the expression-increasing allele is associated with increased methylation. Results for a “secondary CpG” are shown below (descending) in green, for which the expression-increasing allele is associated with decreased methylation. Panel A: eSNP: 10:91165833, expression probe: ILMN_1696654, 1^st^ CpG: cg27582166, 2^nd^ CpG: cg13172359. Panel B: eSNP: 6:37669641, expression probe: ILMN_1720595, 1^st^ CpG: cg26720545, 2^nd^ CpG: cg26129310. Panel C: eSNP: 3:45044878, expression probe: ILMN_2055477, 1^st^ CpG: cg25593573, 2^nd^ CpG: cg06117855.

### Evidence of Shared SNP-by-Sex interaction

Because co-localized eQTL-mQTL pairs are likely to share a biological mechanism, they may also share interactions with factors that modify the effect of an SNP on gene regulation. We tested SNP-by-sex and SNP-by-age interaction for all 519 co-localized eQTLs showing evidence of mediation and/or partial correlation (P<0.05). Only 4 and 2 eQTLs showed evidence of interaction with age and sex, respectively, after Bonferroni correction. Among the four eQTLs showing interaction with age, one (chr18:47043463, P=9×10^−8^ for interaction with age) showed a significant SNP-by-age interaction with respect to the CpG influenced by the co-localized mQTL (P=0.03) (**Figure 8**). The minor allele at this SNP (T) was associated decreased expression of RPL17-C18ORF32 (a read-through transcript) and increased methylation at cg02563971 (assigned to C18ORF32 by Illumina), and both effects were stronger among older participants. Results were similar when analyzing transformed or non-transformed expression and methylation data (**Supplementary Figure 4**). Among the 5 Bonferroni-significant SNP-age interactions for mQTLs, only one (11:118940479 and cg23878202, P=7×10^−7^ for interaction) showed significant SNP-age interaction for the co-localized eQTL (VPS11).

**Figure 8.**
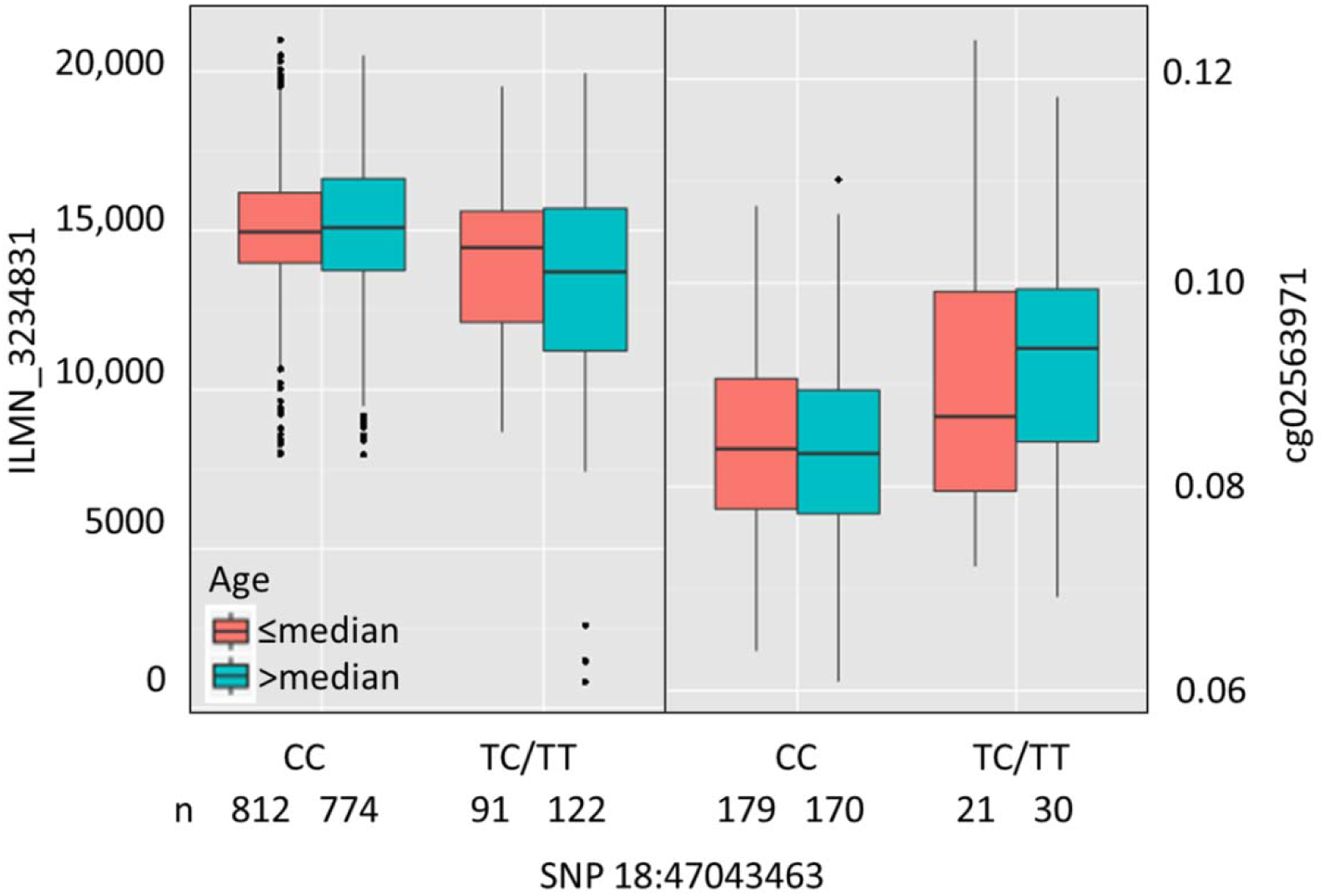
A SNP showing interaction with age in relation to both local gene expression (left) and local DNA methylation (right). Expression probe is ILMN3234831 (RPL17-C18ORF32). The CpG (cg02563971) assigned by Illumina to C18orf32. The P-values for interaction are 9×10^−8^ and 0.03 for the eQTL (n=992) and mQTL respectively (n=337).

## Discussion

In this work, we have described the extent to which peripheral blood eQTLs and mQTLs share common causal variants using data from a Bangladeshi cohort. We identified a set of 3,695 eQTL-mQTL pairs likely to share a CCV, and we used partial correlation analysis and mediation analysis to assess the evidence these pairs of expression and methylation traits are causally related to one another, sharing a common biological mechanism. Among the 3,695 co-localized pairs, we found such evidence for 519 pairs, with mediation and partial correlation analysis showing highly consistent results. Our results demonstrate widespread co-localization of eQTLs and mQTLs in the human genome and shared biological mechanisms. The approach taken here can be extended to other types of cellular/molecular QTLs (e.g., SNPs affecting chromatin features, protein abundance, etc.) in order to enhance our understanding of the cascade of regulatory mechanisms by which SNPs can affect gene expression and function, which is critical for understanding how SNPs affect human disease.

The SNP underlying each co-localized eQTL-mQTL pair tended to have opposite effects on expression and methylation, consistent with the view that hypo-methylation near the promoter and the transcription start site reflects accessible chromatin and active transcription. However, recent work suggests new scenarios in which DNA methylation can also create new binding sites for transcription factors, potentially leading to alternative binding sites in the presence of high levels of methylation ^22^. This is an interesting possibility, considering a subset of our co-localized eQTL-mQTL pairs appear to a) affect expression and methylation in the same direction and/or b) affect multiple CpG sites in opposite directions, suggesting a more nuanced relationship between DNA methylation and local gene expression.

Several prior studies have attempted to assess causal relationships among expression and methylation features that are associated with a common SNP ^14–17^; however, this is the largest such study to date and the first use of co-localization methods to identify relevant eQTL-mQTL pairs. Colocalization is critical for selecting eQTL-mQTL pairs for analyses focused on understanding causal relationships, as such analysis make the implicit assumption that eQTLs and mQTLs share common causal variant. The LD between the lead SNPs for the two QTLs is predictive of the probability of a CCV, but co-localization analysis allows for quantification of the uncertainty regarding the probability that an eQTL and mQTL share a CCV. For a specific mQTL, choosing a CpG remains a challenge, as an mQTL can be associated with increased methylation at one CpG and decreased expression at another, and both can co-localize with an eQTL.

We focus on two approaches for examining evidence that co-localized eQTL-mQTL pairs represent causal SNPs that effect both expression and methylation along a common causal pathway. The first method, partial correlation analysis, detects correlation between expression and methylation that is independent of the regulatory SNP (i.e., residual correlation after adjusting both phenotypes for the SNP). Lack of correlation after adjustment suggests there is not a causal relationship between methylation and expression, as correlation is purely driven by SNP effects ^18^. The second method, mediation analysis, is a test for shared phenotypic variance amongst the SNP, transcript, and methylation. Mediation analysis can also be conceived of as a test for attenuation of the SNP-phenotype relationship after adjusting for a potential mediator. Mediation can be viewed as a more stringent test than partial correlation, as the presence of mediation implies non-zero partial correlation. For both of these methods, we must keep in mind that there are limitations for all tests used to assess evidence of causality; and these tests cannot be used as definitive evidence of causality for any given eQTL-mQTL pair. For example, for some pairs there could exist hidden confounders that are not well captured by the principal components variables we adjust for, and the presence of mediator-outcome confounding can introduce bias into mediation analyses ^23^. In addition, while our Sobel P-values tend to favor the SME model over the SEM, we cannot determine the *direction* of causality for any given pair of expression and methylation traits that appear to be causally related to one another. In other words, for any given pair, it is possible that a) hyper-methylation near the transcription start site makes DNA less accessible for transcription factor binding or b) binding site polymorphisms affect transcription initiation which then in turn affects chromatin structure, including DNA methylation.

Less than half of our eQTLs co-localize with an mQTLs, and several factors may contribute to this observation. First,while co-localization analysis does not require the assumption that there is a single causal variant, it does require the assumption that all causal variants are shared. Thus, it is possible that the presence of a non-shared causal variant near a shared causal variant may lead one to the incorrect conclusion regarding co-localization of the shared variant. Second, we are likely underpowered to detect co-localization when the eQTL and/or eQTL associations are quite weak, as the probability of a CCV clearly depends on the strength of the association. Third, the probability of CCV was systematically lower in high LD regions, making it less likely to detect true co-localization in such regions. Fourth, the RNA and DNA samples used for expression and methylation measurements was not obtained from identical populations of white blood cells (mononuclear cells vs whole blood, respectively). PBMCs (monocytes, T lymphocytes, B lymphocytes) account for ~35% of peripheral white blood cells; thus, the remaining 65% of peripheral white blood cells (neutrophils, basophils and eosinophils) are represented in our DNA methylation data but not in our expression data. Thus, for co-localized eQTL-mQTL pairs that are specific to PMBC subtypes, the mQTL signal may be weak in our data due to the presence of the many cell types in whole blood that are not PBMCs. Lastly, it may be that only a subset eQTLs that impact DNA methylation. In LCLs, for example, it has been reported that only 10–20% of eQTLs are also mQTLs^3^. eQTL mechanisms that would not necessarily involve local epigenetic alterations include effects on mRNA processing or mRNA stability^18^.

Among our 3,695 potentially co-localized pairs, <20% showed strong evidence of mediation and/or partial correlation, and this apparent discrepancy is likely due, at least in part, to several factors that reduce statistical power. First, our mediation and partial correlation analyses are likely underpowered for many of these tests, which require participants with both expression and DNA methylation data. We have only 316 such individuals. In light of the strong association we observe between the strength of the eQTL and mQTL associations and the P-values for our tests of mediation and partial correlation, power is likely to be low for many tests. Second, the cell type issue above will also reduce power to detect mediation and partial correlation, as we are analyzing a mixture of cell types in the presence of cell-type-specific QTLs. The proportion of eQTLs that strongly co-vary with mQTLs in this dataset may be lower than would be observed in a study of similar size focused on a specific cell type, as our methylation measures capture variation in methylation attributable to many different cell types. Third, considering all transcripts and CpGs are imperfect measures, and the CpG we select for analysis is a proxy for some underlying epigenetic state, our power is likely reduced by measurement error. In fact, much of the mediation evidence we detect is “partial mediation” (i.e., mediation proportion <1), and we have shown that this is expected when full mediation is present, but the mediation measure is error-prone^21^.

Additional research is needed to characterize the extent to which co-localized QTLs share interactions with age, sex, or environmental factors (G×E). Sharing GxE would often be expected when expression and methylation causally related, thus providing further evidence for a shared biological mechanism. We provide evidence for one such interaction, and understanding such interactions across multiple molecular phenotypes can further elucidate mechanisms of gene regulation. Future studies should also considering developing more detailed guidance on how to set priors for co-localization analysis, as results are clearly sensitive to choice of priors. In this work we follow the developer's guidance for setting reasonable priors. Lastly, future studies should develop methods for combining data on multiple CpGs to characterize the effects of SNPs on local methylation and chromatin structure. This is important as most mQTLs are associated with multiple CpGs, sometimes in opposing directions.

## Methods

### Study Population

Subjects included in this work were participants in the Bangladesh Vitamin E and Selenium Trial (BEST)^24^. BEST is a randomized chemoprevention trial evaluating the long-term effects of vitamin E and selenium supplementation on non-melanoma skin cancer risk among 7,000 individuals with arsenic-related skin lesions living in seven sub-districts in Bangladesh. Participants included in this work are a subset of BEST participants from Araihazar that have available data on genome-wide SNPs and array-based expression and DNA methylation measures (described below).

### Genotyping, Imputation, and Quality Control

DNA extraction for genotyping was carried out from the whole blood using the QIAamp 96 DNA Blood Kit (cat # 51161) from Qiagen, Valencia, USA. Concentration and quality of all extracted DNA were assessed using Nanodrop 1000. As starting material, 250 ng of DNA was used on the Illumina Infinium HD SNP array according to llluminàs protocol. Samples were processed on HumanCytoSNP-12 v2.1 chips with 299,140 markers and read on the BeadArray Reader. Image data was processed in BeadStudio software to generate genotype calls.

Quality control was conducted as described previously for a larger sample of 5,499 individuals typed for 299,140 SNPs ^25,26^. We removed DNA samples w samples with call rates <97% (n = 13), gender mismatches (n = 79), as well as technical duplicates (n=53). We removed and SNPs that were poorly called (<90%) or monomorphic (n = 38,753), and then removed SNPs with call rates <95% (n = 1,045) or HWE p-values<10^−10^ (n = 1,045). This QC resulted in 5,354 individuals with high-quality genotype data for 257,747 SNPs. The MaCH software^27^ was used to conduct genotype imputation using 1,000 genomes reference haplotypes (including South Asian populations). Only high-quality imputed SNPs (imputation r^2^>0.5) with SNPs with MAF>0.05 were included in this analysis. A subset 1,329 unrelated individuals with available data on array-based expression and DNA methylation measures was used for this project. Only autosomal SNPs were included in this analysis.

### DNA methylation

DNA was extracted from whole blood using DNeasy Blood kits (Qiagen, Valencia, CA, USA). Bisulfite conversion was performed using the EZ DNA methylation Kit (Zymo Research, Irvine, CA, USA). For each sample, 500 ng of bisulfite-converted DNA was applied to the Illumina HumanMethylation 450K BeadChip kit (Illumina, San Diego, CA, USA) according to the manufacturer’s protocol, enabling interrogation of 482,421 CpG sites and 3,091 non-CpG sites per sample. This array contains an average of 17 CpG sites per gene, distributed across the promoter, 5̀ UTR, first exon, gene body and 3́ UTR, covering 99% of RefSeq genes.

Methylation status at each CpG is expressed as a β value that can range from 0 (unmethylated) to 1 (completely methylated). Data were quantile normalized. Among the 413 participants, we excluded 6 samples for which the reported sex of the participant did not correspond with predicted sex based on methylation patterns of the X and Y chromosomes, and 7 samples with > 5% of CpGs either containing missing values or having p for detection > 0.05. This resulted in 400 samples with quality methylation data. We removed probes mapping to multiple locations (41,937) and probes with SNPs (20,869) according to Price et al. Individual β values with a *p* for detection > 0.05 were set to missing, and we excluded probes if >10% of beta values were missing (1,636). We also excluded probes on the X (11,232) and Y (416) chromosomes, probes with missing chromosome data (mostly control probes, 65), and probes with > 10% missing data across samples (1,932); this resulted in a total of 423,604 probes available for analysis. β values were logit transformed and adjusted for batch variability using ComBat software^28^. Based on 11 samples run in duplicate across two different plates in these experiments, the average inter-assay Spearman correlation coefficient was 0.987 (range, 0.974–0.993). There were six individuals excluded whose self-reported sex did not match their sex based on methylation data, resulting in 407 individuals with high quality methylation data.

### Gene Expression

RNA was extracted from PBMCs, preserved in buffer RLT, and stored at −86°C using RNeasy Micro Kit (cat# 74004) from Qiagen, Valencia, USA. Concentration and quality of RNA samples were assessed on Nanodrop 1000. cRNA synthesis was done from 250 ng of RNA using Illumina TotalPrep 96 RNA Amplification kit. As recommended by Illumina we used 750 ng of cRNA on HumanHT-12-v4 for gene expression. Expression data were quantile normalized and log_2_ transformed. The chip contains a total of 47,231 probes covering 31,335 genes. There were 1,825 unique individuals in both expression data and SNP data. For the vast majority of participants, between 30% and 47% of probes had detection P values <0.05. However, 26 individuals had>30% of probes with detection p-value <0.05, and these outlying individuals were excluded from the analysis, leaving an analysis sample size of 1,799. For this analysis, no probes were excluded based on detection P-values.

### Eligibility for analyses

The participants and workflow are described in **Figure 1** and **Supplementary Figure 1**. Participants included in eQTL analyses included 992 participants with available SNP data and expression data who were unrelated to other participants based on an estimated coefficient of relationship <0.08. Participants included in mQTL analyses included 337 participants with available SNP data and DNA methylation data who were unrelated to other participants based on an estimated kinship coefficient of <0.08. These samples used for eQTL and mQTL analyses were entirely independent (i.e., non-overlapping participants), which is a requirement for using co-localization methods^19^. Among the 337 participants included in mQTL analyses, 316 of these participants also had expression data (which was not used for eQTL analyses), and these 316 participants were used for mediation analyses, Bayesian network analyses, and partial correlation analyses.

### eQTL and mQTL analyses

Prior to analysis, expression values were log transformed and methylation beta values were logit-transformed and adjusted for potential batch/chip effects. Linear regression implemented in the matrix-eQTL software package^29^ was used to conduct genome-wide *cis*-eQTL and and *cis*-mQTL analyses. *Cis* associations were tested for SNPs and probes <500 Mb apart. For both the *cis-eQTL* and mQTL analyses, we used an FDR of 0.01 to report significant associations (using the Benjamini and Hochberg method). In addition to adjusting for age and sex, we included 80 expression PCs in our eQTL analyses and 10 methylation PCs in our mQTL analyses, and these were selected to maximize the number of cis signals detected ^21^. Lead eSNPs and mSNPs for each eGene and mCpG, respectively, were defined as the SNP with the smallest P-value.

### Identification of eQTL-mQTL pairs that potentially share a common causal variant

Our workflow for identifying co-localized eQTL/mQTL pairs (sharing a common causal variant) is shown in **Supplementary Figure 1**. For our eQTL results, we first restricted to lead SNPs for each eProbe. Using the mQTL results, we then identified CpGs that were also associated with a lead eSNP. Because clusters of CpGs are often correlated and influenced by the same cis-variation^30^, we pruned our list of CpG probes to reduce this redundancy. We pruned by first identifying CpGs that were associated with the same SNP, and kept only the CpG whose lead mSNP had the highest LD with a lead eSNP. We required each expression probe to be in a pair with only one CpG, the CpG whose lead mSNP was in the strongest LD with the expression probe’s lead eSNP. This workflow resulted in 7,656 eProbe-CpG pairs showing association with a common SNP and able to be tested for colocaliation.

### Co-localization analysis

To assess the probability that cis-mQTLs and cis-eQTLs residing in the same genomic location were due to the same (single) causal variant, we applied a Bayesian test for co-localization ^19^ to all co-occurring eQTL-mQTL pairs, in order to estimate the probability that each QTL pair were due to the same causal variant. The Bayesian co-localization requires specifying a prior probability for a SNP being associated with trait 1 (p_1_), trait 2 (p_2_), and both traits (p_12_). We used a prior probability (p) of 10^−4^ for a SNP being associated with the expression trait (p_1_=10^−4^) and a SNPs being associated with a methylation trait (p_2_=10^−4^), as recommended by the developers. Following the developer's approach ^19^, we varied the value of p_12_ (10^−5^, 10^−6^, and 10^−7^) in order to evaluate the sensitivity of the results to the prior. Our p_12_ values of 10^−5^, 10^−6^, and 10^−7^ are interpreted as 1 in 10, 1 in 100, and 1 in 1000 probability that a SNP associated with expression is also associated with methylation (or vice versa).

### Partial Correlation Analysis

Using our set of 3,695 potentially co-localized eQTL-mQTL pairs, we used data on 316 genotyped individuals with both expression and methylation data to conduct partial correlation analysis^18^. We first calculated the Pearson correlation coefficient between the expression probe and the methylation probe (both adjusted for expression and methylation PCs, respectively, as described above). We then regressed both the methylation probe and the expression probe on the lead SNP, and took the residuals from these regressions to obtain expression and methylation values that lack the phenotypic variance due to the effect of the SNP. We then compare the correlation coefficient before SNP adjustment vs. after SNP adjustment to test the independence of the eSNP on methylation/expression.

### Mediation Analysis Methods

Using our set of 3,695 potentially co-localized eQTL-mQTL pairs, we used data on 316 genotyped individuals with both expression and methylation data to conduct tests of mediation for two hypothesized pathways: 1) SNP -> methylation -> Expression (“SME”) and 2) SNP -> Expression -> methylation (“SEM”). Mediation analysis was conducted as follows: For all lead eSNP, the cis-eQTL association was re-estimated, adjusting for methylation of the CpG (and vice versa). The difference between the beta coefficients before and after adjustment for the *cis* probe was expressed as the “proportion of the total effect that is mediated” (i.e., % mediation), calculated as (β_unadj_ – β_adj_)/ β_unadj_^31^, with β_unadj_ and β_adj_ known as the *total effect* the *direct effect*, respectively. All regressions were adjusted for expression and methylation PCs. The Sobel P-value for mediation^32^ was calculated by first estimating the cis-eQTL association adjusting for methylation (and vice versa):

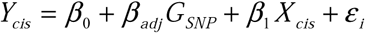

We then estimated the eSNP’s association with the potentially mediator:

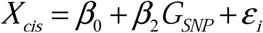

The P-value was then estimated by comparing this following t statistic to a normal distribution:

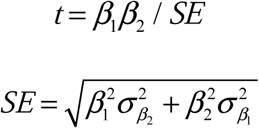

where SE is the pooled standard error term calculated from the above beta coefficients and their variances, β_1_ β_2_ is often referred to as the *indirect effect*.

### Mediation analysis of simulated data

Using data on a bi-allelic SNP (G) for 316 participants (same sample sizes as our analyses), we simulated data on a molecular phenotype (X) as a randomly generated standard normal variable with a linear effect exerted by the SNP

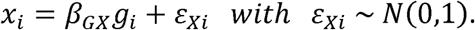

X served as a mediator for the effect of the SNP on a second molecular phenotype (Y), which was generated as a standard normal variable with a linear effect exerted by the (X).

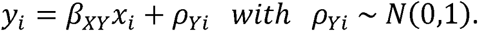

The variance in the mediator (X) explained by the SNP was varied from 0.01 to 0.75. The magnitude of the effect of the mediator on the second molecular phenotype (β_XY_) was varied from 0.01 to 0.75. We then used mediation analysis methods described in the section above to obtain a Sobel P-value and an estimate of the % mediation. These analyses were conducted in two ways: using X as the mediator and using Y as the mediator.

## Acknowledgements

We would like to thank all BEST study participants and research staff. This work was supported by National Institutes of Health grants R21 ES024834 (B.P. and M.A.), ROI ES020506 (B.P.), ROI ES023834 (B.P.), and ROI GM108711 (L.C.)

## Author contributions

B.L.P., L.C., and H.A developed the general research question. B.L.P. developed the analysis approach, supervised the analyses, and drafted the manuscript. L.T. conducted all statistical analyses. H.A. established BEST and led the collection of all bio-specimens and generation of SNP, expression, and methylation data. M.K. and J.F supervised and conducted DNA and RNA extraction and the generation of SNP, expression, and methylation data. L.C. provided assistance with statistical aspects of the work and manuscript preparation. M.A., M.R-Z., G.S., M.T.I, H.S., T.I., M.R, and M.Y. contributed to BEST study design and data collection. All authors reviewed and edited the manuscript.

